# Mechanical Fidelity of Fixed-Cell Traction Force Microscopy

**DOI:** 10.64898/2026.11.08.765148

**Authors:** Armina Mortazavi, Jianfei Jiang, Benedikt Sabass

## Abstract

Cellular traction forces contribute to biological processes including cell migration and tissue morphogenesis. Traction force microscopy (TFM) is widely used to quantify these forces in living cells, but it remains unclear how well measurements after fixation reproduce the live-cell state. Here, we systematically evaluated fixation-induced changes by paired comparison of the same cells before and after fixation using different fixation protocols and substrate stiffnesses. Fixation largely preserved the spatial organization and direction of substrate displacements and reconstructed traction fields, whereas displacement, traction, and strain-energy magnitudes showed fixation-dependent changes. Among the protocols tested, glutaraldehyde (GA) provided the closest agreement with live-cell measurements, retaining up to 99% of displacement and traction magnitudes and strain energy. Paraformaldehyde-based protocols caused larger reductions in these quantities while largely preserving their spatial organization. These results show that fixed-cell TFM, particularly with GA fixation under the conditions tested here, can provide measurements close to the live-cell state and offers a practical approach for combining TFM with post-fixation fluorescence imaging in a multi-well workflow.

## Introduction

Cells generate and transmit mechanical forces to their microenvironment, contributing to adhesion, migration, differentiation, and morphogenesis^1,2^. Traction force microscopy (TFM) is a widely used approach to quantify cell-generated stresses extracellularly^3–5^. In standard TFM, cells are cultured on soft elastic substrates embedded with fluorescent fiducial beads, and the resulting substrate deformations are tracked relative to a reference image. By combining bead displacement measurements with substrate elasticity, traction forces can be reconstructed, allowing detailed mapping of cellular force patterns^6^. Usually, TFM is performed on live cells to capture the dynamics of traction generation and remodeling. However, live-cell imaging is generally incompatible with conventional immunolabeling and imaging molecular-scale structures is further challenged by phototoxicity, cell motion, limited acquisition times, and the optical constraints due to the live-cell microscopy setup ^7^.

Fixation is widely used to preserve cellular structure and enable high-resolution microscopy by stabilizing biological samples in a defined state^8,9^. Compared to live-cell imaging, fixed samples offer improved reproducibility, compatibility with fluorescence-based techniques, and elimination of motion artifacts, facilitating detailed structural and molecular analysis^10–12^. However, fixation halts dynamic cellular processes and can introduce artifacts such as cell shrinkage, membrane blebbing, cytoskeletal collapse, and redistribution or loss of soluble proteins, with the severity depending strongly on the fixative used^13–17^.

Mechanical properties of live cells and fixed cells have already been compared with different methods. Atomic force microscopy (AFM) indentation revealed that paraformaldehyde fixation can increase apparent stiffness several-fold relative to live cells, consistent with enhanced cross-linking of cytoskeletal structures^18–20^. Using optical tweezers indentation and microfluidic constriction assays, it was demonstrated that chemical fixation preserves overall cell morphology while significantly increasing cellular rigidity and altering dynamic viscoelastic and active responses^21,22^. Collectively, these studies emphasize that fixation can stabilize structural features while altering quantitative mechanical properties, highlighting the importance of assay-specific evaluation.

Cell fixation is increasingly used in combination with TFM, either as a substitute for live-cell measurements in high-throughput reference-free TFM or by fixing cells after performing live-cell TFM^23–26^. In these studies, characteristic deformation patterns are often assumed to be sufficient to eliminate the need for a reference state obtained through cell detachment methods such as trypsinization or surfactant treatments (e.g., SDS or Tween)^27^. However, it remains unclear how accurately fixed-cell TFM reproduces live-cell traction measurements and how strongly the results depend on the fixation protocol. In the context of TFM, fixation can in principle provide several advantages. First, it immobilizes cells and prevents cytoskeletal remodeling during extended imaging, thereby stabilizing substrate displacement fields for long acquisitions. This is particularly advantageous in combination with single-molecule localization microscopy and other super-resolution techniques that require thousands of frames^28,29^. Second, fixation allows direct correlation of traction measurements with immunofluorescence staining, enabling molecular-level interpretation of force patterns. Third, fixed-cell TFM could facilitate high-throughput or retrospective analyses, as samples can be prepared and imaged independently of cell viability constraints^25^. Despite these advantages, a key unresolved question is whether the substrate displacements measured after fixation faithfully represent those measured in live-cell conditions, and how experimental conditions influence this fidelity.

To assess how faithfully fixed-cell measurements reproduce live-cell traction forces and to determine how different fixation methods influence the preservation of traction forces, we systematically compared substrate displacement field, traction, and energy in live and fixed cells across multiple fixation methods and substrate stiffnesses. Broadly, the results demonstrate that fixed-cell TFM can be implemented in comparative studies and provide a basis for combining traction measurements with advanced fluorescence imaging.

## Results

### A multi-well imaging workflow for reliable live-fixed registration

To enable paired live– and fixed-cell TFM measurements with reliable spatial registration, we established an imaging workflow using multi-well chambers and motorized stage acquisition. The rectangular geometry of the wells minimized rotational repositioning errors and enabled consistent relocation of identical fields of view across sequential experimental steps, including reference, live, and post-fixation imaging (Fig. 1a). Reference images of the undeformed substrates containing randomly distributed beads were acquired prior to cell seeding, providing a baseline for subsequent deformation analysis. Imaging regions were optimized to ensure sufficient sampling of isolated single cells while maintaining parallel acquisition across multiple wells. Using a 40× objective, nine adjacent fields of view were acquired in a 3×3 tiled configuration per well and computationally stitched, generating composite imaging areas of approximately 650 µm × 650 µm (Supplementary Fig. 1). This procedure enabled rapid sequential imaging while preserving spatial continuity and registration accuracy between tiled regions (Fig. 1b). Importantly, we did not require substrates with regular spatial patterns for consistent relocation of identical fields of view across experimental stages through stored motorized stage coordinates. Following cell seeding and incubation, live-cell brightfield and fluorescence images were acquired from predefined regions to record cell morphology and fiducial positions. Samples were then fixed, and identical field of views were imaged again under matched acquisition conditions. In subsequent analysis, we compared live– and fixed-cell deformation fields to quantify fixation-induced changes in traction forces and derived energetic metrics (Fig. 1c). All imaging steps (reference, live, and post-fixation) were performed strictly at room temperature, and image registration was corrected using cell-free background areas.

**Fig 1.**
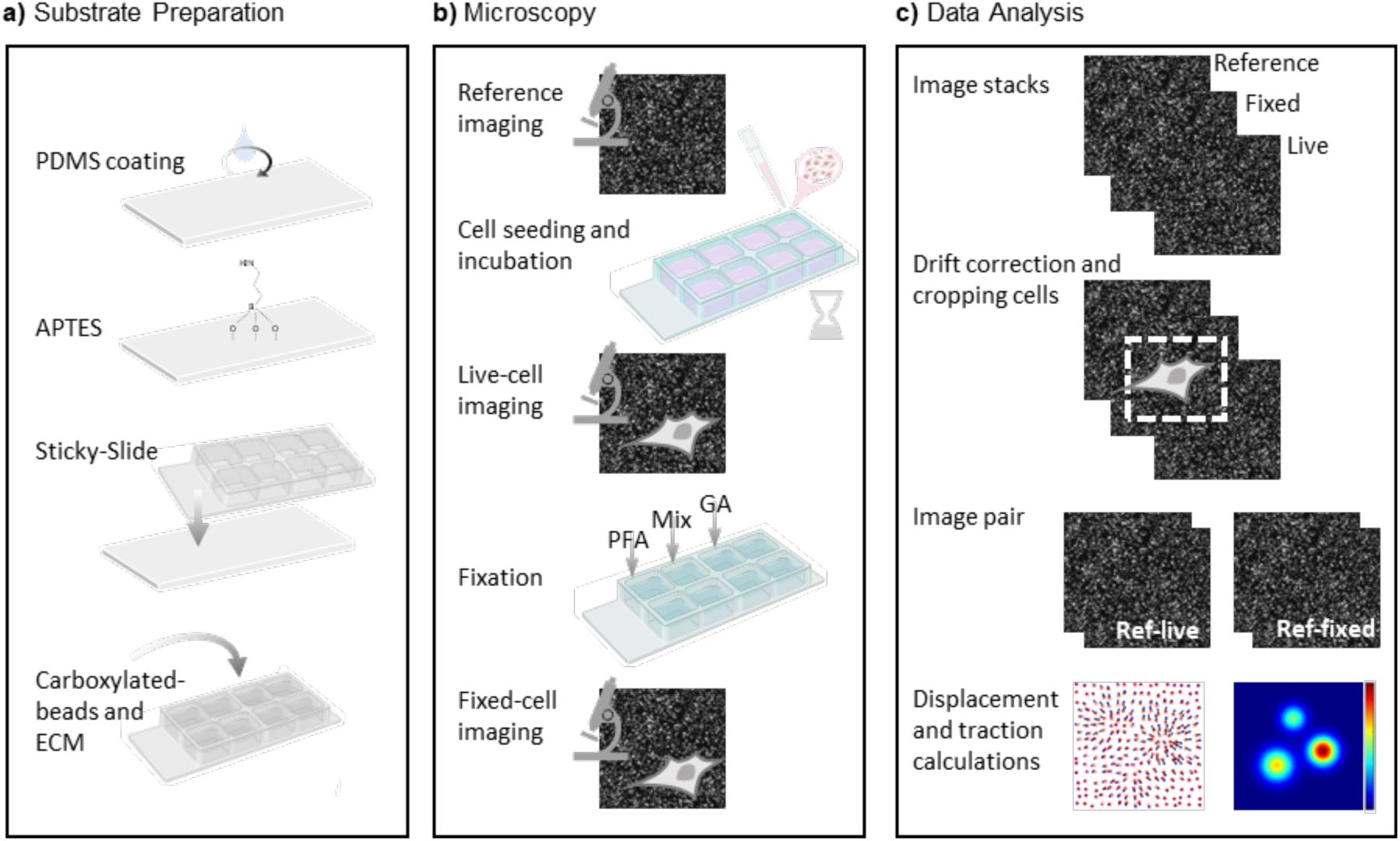
Experimental workflow, imaging procedure, and data processing. (a) Substrate preparation: Glass coverslips were coated with polydimethylsiloxane (PDMS) and functionalized with aminopropyltriethoxysilane (APTES), then assembled into 8-well chamber slides. Substrates were subsequently functionalized with fluorescent beads and coated with extracellular matrix (ECM) proteins to enable traction force microscopy measurements. (b) Microscopy and imaging workflow: A reference image of the substrate containing beads was first acquired in the absence of cells. Cells were then seeded and incubated to allow adhesion and force generation. Imaging was performed to capture both bead displacement fields and corresponding cell morphology at identical positions. Samples were subsequently fixed using paraformaldehyde (PFA), a paraformaldehyde-glutaraldehyde mixture (Mix), or glutaraldehyde (GA), followed by post-fixation imaging of the same fields of view. (c) Data analysis pipeline: Image stacks from reference, live, and fixed conditions were aligned to correct for stage drift. Regions containing cells were identified and cropped. Matched image pairs (live– reference and fixed–reference) were then used for traction force reconstruction and downstream quantitative analysis.

### Substrate displacements maintain their spatial structure during fixation

To assess how fixation affects substrate deformation, we compared displacement directions and magnitudes for the same cells before and after fixation on substrates of different stiffnesses (Fig. 2a–c). Directional cosine values between matched displacement vectors were close to unity (0.97–0.98; Fig. 2d), with no significant effect of fixation method (*p* = 0.435; Supplementary Table 1). Spatial correlations between live– and fixed-cell displacement-vector components were similarly high (0.94–0.96; Fig. 2e) and did not differ significantly among fixation methods (*p* = 0.839). Thus, fixation largely preserved the direction and spatial organization of substrate displacements.

**Fig. 2:**
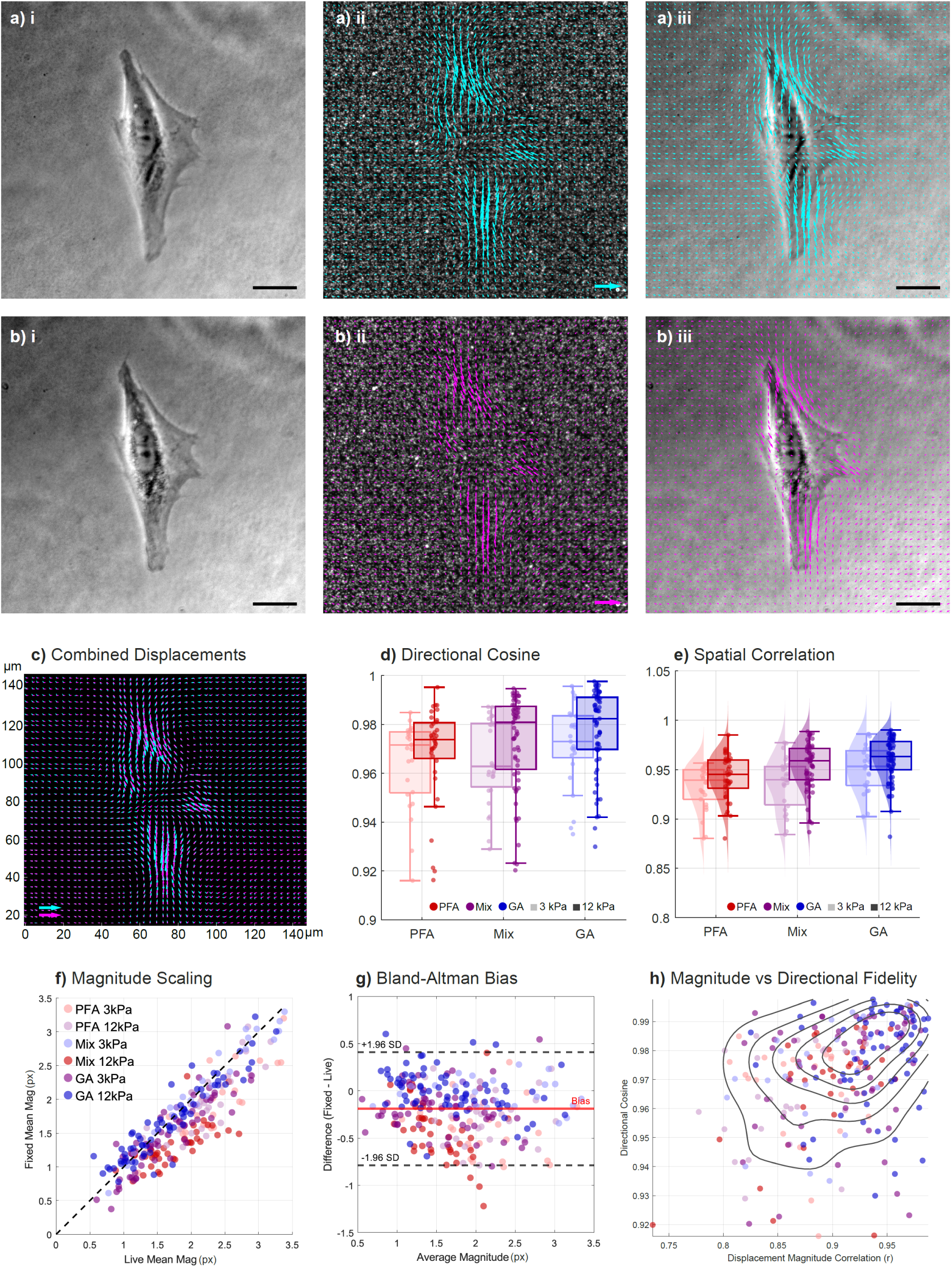
**Comparison of substrate displacements caused by cells before and after fixation**. (a) Live-cell imaging. (i) Representative bright-field image of a live human foreskin fibroblast (HFF) cell, seeded on PDMS substrate with Young’s modulus of 12 kPa (scale bar = 20 µm). (ii) fluorescent beads with overlaid displacement vectors. Vectors are magnified 5× for visualization; the reference arrow corresponds to a true displacement of 10 µm. (iii) Superimposed view of the cell with displacement field, showing spatial organization of traction forces relative to cell morphology. (b) Fixed cells: (i) Bright-field image of an HFF cell after fixation with a mix of paraformaldehyde (PFA) and glutaraldehyde (GA); (scale bar = 20 µm). (ii) fluorescent beads with overlaid displacement vectors (same 5× magnification and 10 µm reference scale as in (a)). (iii) Superimposed view of the fixed cell with displacement field. (c) Combined comparison: Overlay of live (cyan) and fixed (magenta) displacement fields showing preserved spatial structure and directionality across conditions, plotted with identical vector scaling. (d) Directional cosine: Quantification of directional agreement between live and fixed displacement vectors, computed as the cosine similarity between corresponding vector pairs. Values close to 1 indicate strong preservation of displacement direction after fixation. (e) Spatial correlation. Pearson correlation coefficient between spatially matched displacement-vector components (*u_x_*, *u_y_*) across all grid points (*r*_spatial_ = corr(**u**_live_, **u**_fixed_)), quantifying the preservation of spatial deformation patterns across conditions. corresponding displacement-vector components in live and fixed fields, quantifying preservation of spatial deformation patterns across conditions. High values indicate strong conservation of the displacement pattern. (f) Magnitude scaling. Scatter plot comparing live and fixed displacement magnitudes for all cells. The dashed unity line represents perfect agreement between live and fixed measurements. Data points falling below the unity line indicate underestimation of displacement magnitude after fixation. (g) Bland–Altman analysis: Comparison of live– and fixed-cell displacement magnitudes, plotted as the difference versus the mean of paired measurements. The plot quantifies systematic bias and limits of agreement across the measurement range. The solid line indicates the mean bias, and the dashed lines represent the limits of agreement (±1.96 SD), revealing a systematic reduction in displacement magnitude after fixation. (h) Spatial magnitude correlation versus directional fidelity: Relationship between displacement magnitude correlation and directional cosine, assessing whether spatial magnitude preservation post-fixation is correlated to directional distortions. Weak correlation indicates independent mechanisms governing magnitude and direction fidelity. A total of 267 cells from 3 independent experimental preparations were analyzed. Each included both substrate stiffnesses (3 and 12 kPa) and all three fixation protocols (PFA, Mix, and GA). Color coding: Red, purple, and blue denote PFA, Mix, and GA fixation, respectively. Light/translucent shades indicate 3-kPa substrates, whereas darker, saturated shades indicate 12-kPa substrates.

We next compared displacement magnitudes before and after fixation. For each sample, the magnitude of each displacement vector was calculated and averaged over the analyzed field of view (Fig. 2f). The mean displacement magnitudes before and after fixation were strongly correlated (Pearson’s *r* = 0.880), but magnitudes were generally lower after fixation. Mean fixed-to-live displacement ratios were 0.738 ± 0.139 for PFA, 0.846 ± 0.136 for the mixed protocol, and 0.986 ± 0.113 for GA, corresponding to retention of approximately 74%, 85%, and 99%, respectively.

Across all conditions (*n* = 3 wells per condition; *N* = 267 cells), mean displacement magnitude decreased from 2.00 ± 0.83 px in live cells to 1.80 ± 0.74 px after fixation. Bland–Altman analysis showed a mean bias of −0.189 ± 0.305 px, with limits of agreement from −0.787 to 0.409 px (Fig. 2g). The absolute bias was slightly larger on 3-kPa substrates (−0.226 ± 0.293 px) than on 12-kPa substrates (−0.172 ± 0.310 px), consistent with the larger displacements observed on softer substrates. Finally, displacement magnitude correlation and the cosine of the directional deviation were only weakly associated (Pearson’s *r* = 0.358, *P* < 0.001; *R*² = 0.128; Fig. 2h). Thus, fixation largely preserved displacement direction and spatial organization, whereas displacement magnitude was more sensitive to the fixation protocol, with GA providing the highest fidelity.

### The magnitude of the recovered tractions depends on the fixation protocol

We next examined whether the effects of fixation on substrate displacements were also reflected in the reconstructed traction forces. Representative traction maps showed that the spatial distribution of tractions was largely preserved after fixation, whereas traction magnitudes were reduced (Fig. 3a).

**Fig. 3:**
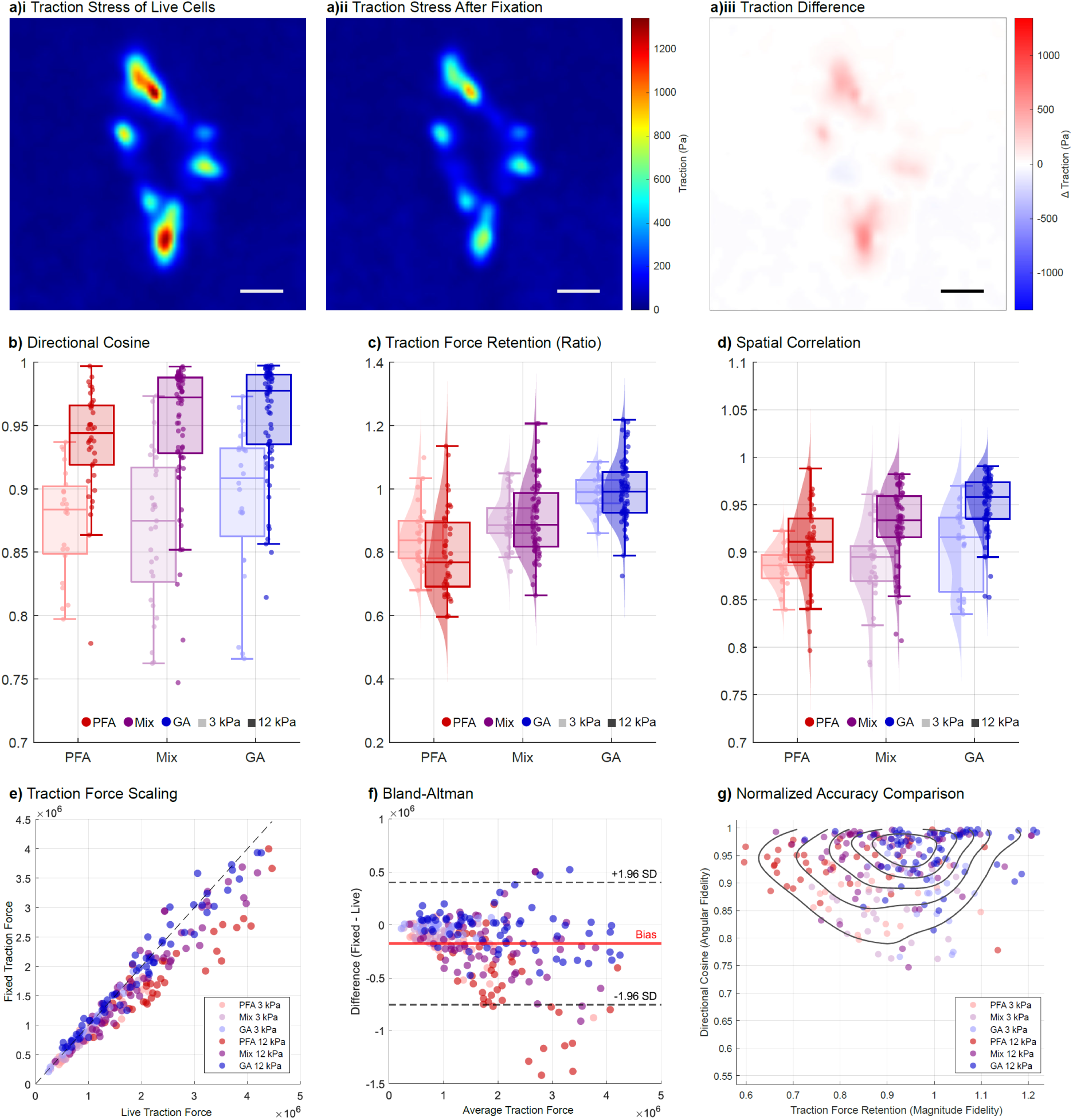
**Traction forces before and after fixation**. (a) Representative traction maps. (i) Traction forces generated by a live HFF cell, seeded on a PDMS substrate with Young’s modulus of 12 kPa. (ii) Corresponding traction map after fixation with a mix of PFA and glutaraldehyde GA (iii) Magnitude of the difference between live– and fixed-cell traction fields, highlighting fixation-induced changes. (b) Directional cosine: Quantification of angular agreement between live and fixed traction vectors. (c) Ratio of fixed– to live-cell traction force, representing preservation of force magnitude across fixation conditions. (d) Spatial correlation: Correlation analysis of spatially matched traction fields between live and fixed states, assessing conservation of global force organization and traction pattern architecture. (e) Traction force scaling comparison: Scatter plot comparing live and fixed total traction forces. The dashed unity line represents perfect agreement. Systematic deviation below the unity line indicates underestimation of force magnitude after fixation. (f) Bland–Altman analysis: Difference-versus-mean comparison of live and fixed traction forces. (g) Normalized accuracy comparison: Relationship between force retention and directional cosine, comparing magnitude fidelity and angular fidelity across fixation methods and substrate stiffnesses. Density contours indicate regions of high sample concentration and illustrate the weak coupling between magnitude preservation and directional accuracy. A total of 267 cells from 3 independent experimental preparations were analyzed. Each included both substrate stiffnesses (3 and 12 kPa) and all three fixation protocols (PFA, Mix and GA). Color coding: Red, purple, and blue denote PFA, Mix, and GA fixation, respectively. Light/translucent shades indicate 3-kPa substrates, whereas darker, saturated shades indicate 12-kPa substrates.

Directional fidelity was quantified as the cosine similarity between live and fixed traction vectors exceeding 20% of the maximum live-cell traction magnitude to exclude spatial regions with low background traction (Fig. 3b). Cosine similarity was high across conditions, ranging from 0.861 ± 0.050 for PFA on 3-kPa substrates to 0.955 ± 0.048 for GA on 12-kPa substrates. Fixation method (*p* = 0.435) and the method × stiffness interaction (*p* = 0.540) had no significant overall effect. However, post-hoc analysis showed higher directional fidelity for GA than for PFA (*P* < 0.05).

To compare traction magnitudes before and after fixation, the magnitude of each traction vector was calculated and averaged over the analyzed field for each sample. Traction magnitude retention was defined as the ratio of the mean traction magnitude after fixation to the corresponding live-cell value. Retention depended significantly on fixation method (*p* = 0.0169), with no significant method × stiffness interaction (*p* = 0.941; Fig. 3c). GA showed the highest retention (0.986 ± 0.053 and 0.992 ± 0.096 for 3 and 12 kPa, respectively), followed by the mixed protocol (0.901 ± 0.077 and 0.906 ± 0.116) and PFA (0.850 ± 0.094 and 0.795 ± 0.135). Spatial correlations between live and fixed traction magnitudes were also high across conditions (Fig. 3d), reaching 0.950 ± 0.030 for GA on 12-kPa substrates.

Live and fixed traction magnitudes were strongly correlated overall (Pearson’s *r* = 0.957; Fig. 3e), although values after fixation were generally lower. Bland–Altman analysis confirmed this negative bias and showed increasing absolute differences at higher traction magnitudes (Fig. 3f). Force retention and directional fidelity were only weakly associated (*r* = 0.087, *P* = 0.164; *R*² = 0.008; Fig. 3g).

Thus, fixation largely preserved the spatial organization and direction of traction forces, whereas traction magnitude depended more strongly on the fixation protocol, with GA providing the closest agreement with live-cell measurements.

### Strain energy after fixation of cells

We finally assessed the effect of fixation on the cellular strain energy stored in the substrate. Strain energy retention was defined as the ratio of strain energy after fixation to the corresponding live-cell value. Fixation resulted in an overall reduction in strain energy (relative difference: −0.114 ± 0.168, *P* < 0.001). Energy retention depended significantly on fixation method (*p* = 0.0221; Fig. 4a,b), with GA showing near-complete retention (99.4 ± 5.5%) and significantly higher retention than PFA (Tukey post-hoc, *p* = 0.0224).

**Fig. 4:**
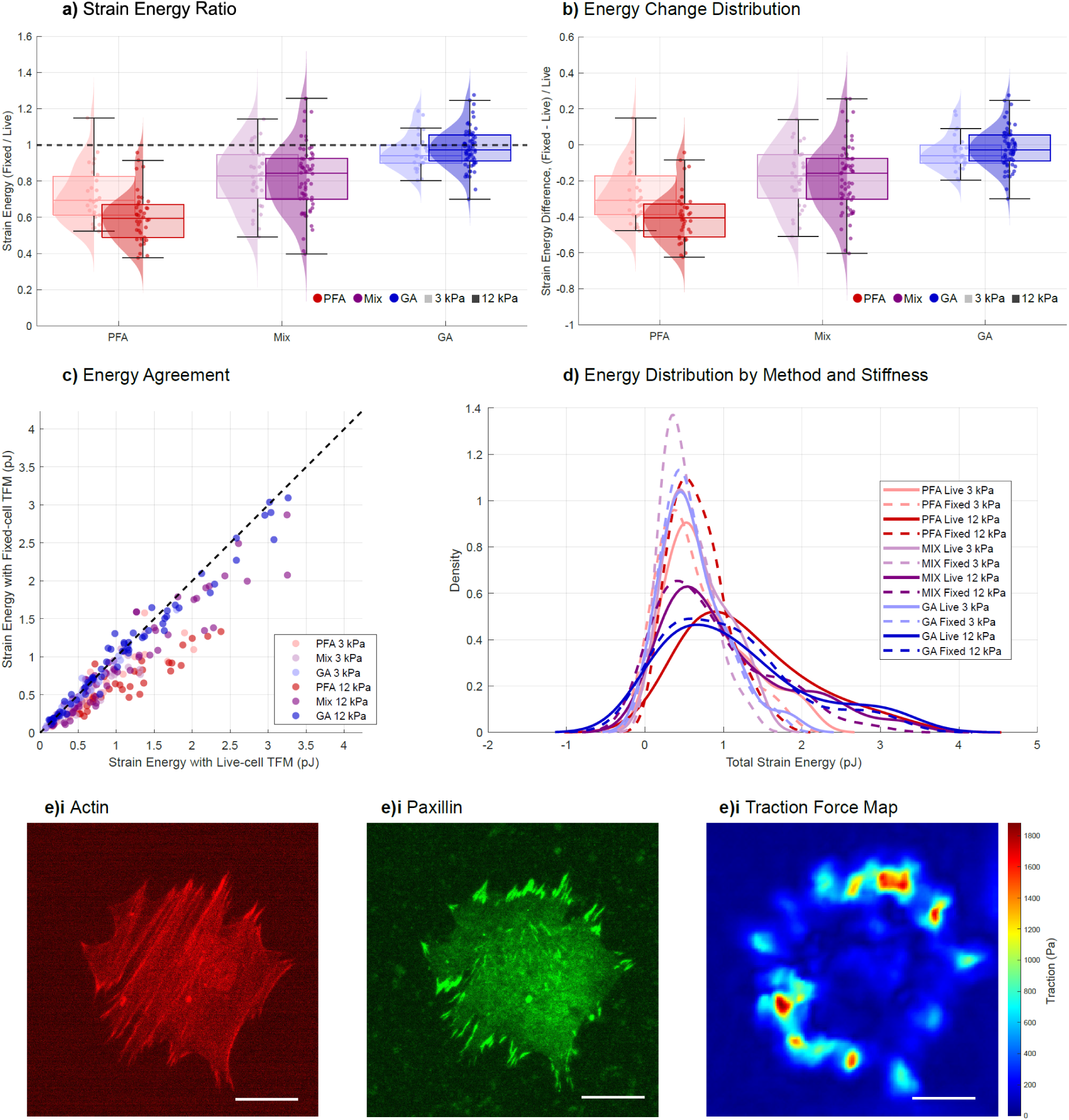
**Energetic analysis and structural characterization of fixed-cell traction preservation**. (a) Distribution energy retention values, calculated as the ratio of fixed to live strain energy across different fixation methods and substrate stiffnesses. (b) Distribution of relative energy differences between fixed and live conditions, illustrating the magnitude and variability of fixation-induced energy loss across samples. (c) Scatter plot comparing live and fixed strain energy measurements. Each dot corresponds to one cell. The dashed unity line represents perfect agreement. (d) Strain-energy distributions by method and substrate stiffness. For data shown in (a)-(d), a total of 267 cells from 3 independent experimental preparations were analyzed. Each included both substrate stiffnesses (3 and 12 kPa) and all three fixation protocols (PFA, Mix and GA). Color coding: Red, purple, and blue denote PFA, Mix, and GA fixation, respectively. Light/translucent shades indicate 3-kPa substrates, whereas darker, saturated shades indicate 12-kPa substrates. (e) Representative example illustrating the cytoskeletal structure and traction forces after fixation on an elastic substrate. The depicted cell was seeded on a PDMS substrate with Young’s modulus of 12 kPa and fixed with PFA. (i) Phalloidin-stained actin cytoskeleton following fixation. (ii) A paxillin stain shows the organization of focal adhesions. (iii) Reconstructed traction force map of the same fixed cell, illustrating the relationship between preserved cytoskeletal architecture, adhesion structure, and traction force generation after fixation. Scale bar: 20 µm.

Live and fixed strain energies were strongly correlated across all measurements (Pearson’s *r* = 0.921, *R*² = 0.848; Fig. 4c). The strain-energy distributions were shifted toward lower values after fixation, with the smallest changes observed for GA (Fig. 4d). Thus, consistent with the displacement and traction analyses, GA provided the closest agreement between live and post-fixation measurements.

### Immunofluorescence staining

Finally, we performed Immunofluorescence staining of F-actin (phalloidin) and paxillin following fixation to assess whether structural cytoskeletal organization and focal adhesion architecture are consistent with the mechanically derived traction patterns, see Fig. 4e for an example. The spatial distribution of paxillin-positive focal adhesions and actin stress fibers showed strong agreement with regions of elevated traction stress in the reconstructed traction force heat maps. High-traction regions co-localized with prominent focal adhesion sites and aligned actin filament bundles, supporting the mechanical validity of the post-fixation traction reconstruction. This correspondence further demonstrates that traction force distributions can be robustly extracted after fixation while preserving biologically meaningful spatial information, and that the reconstructed stress fields reflect underlying subcellular force-generating structures.

## Discussion

Chemical fixation of cells is an essential approach in the biological sciences. In some cases, a lack of alternatives to fixation has even precluded a direct comparison to live-cell controls. Among different methods of fixation, alcohol-based fixatives (e.g., methanol) can cause substantial dehydration and cellular shrinkage, whereas PFA generally preserves overall morphology but may still induce subtle changes in cell volume. GA provides stronger crosslinking, resulting in improved stabilization of ultrastructure, although it can also introduce stiffness-related distortions and increased autofluorescence. In practice, PFA and GA are considered among the most reliable fixatives for preserving cellular architecture, with GA offering superior structural stabilization at the expense of potential chemical and optical artifacts^30,31,17^.

Here, we set out to determine how chemical fixation alters cellular force measurements in TFM, considering effects at all levels, from substrate displacements to traction forces and strain energy. Our results suggest that the effects of chemical fixation on TFM measurements can be separated into two aspects: (i) geometric fidelity, encompassing direction and spatial organization, and (ii) magnitude scaling of displacements and tractions. Overall, we find that careful cell fixation preserves the spatial organization of displacements and traction forces, whereas their magnitudes are consistently reduced, although to varying degrees. These fixation artifacts are strongly dependent on the fixation method, with GA providing the highest fidelity. The different results for PFA and GA fixation possibly result from a combination of their different cross-linking kinetics and osmotic effects during aldehyde penetration. PFA is a small, monofunctional aldehyde that forms cross-links relatively slowly, potentially creating a time window during which active cellular tension relaxes before the cytoskeleton is immobilized. In addition, PFA solutions, particularly without membrane-stabilizing agents or carefully balanced tonicity, can induce transient osmotic swelling prior to complete cross-linking, which could mechanically perturb focal adhesions independent of cross-linking speed. GA, as a larger, bifunctional aldehyde, forms cross-links more rapidly, which may more effectively immobilize the cytoskeleton before either tension relaxation or osmotic effects can act. Furthermore, prior to fixation, live-cell medium was replaced with standard Dulbecco’s phosphate-buffered saline (DPBS, lacking Ca^2+^ and Mg^2+^). Because divalent cations stabilize integrin binding and cytoskeletal structural integrity, transient incubation in cation-free buffer could induce minor, rapid tension relaxation independently of fixative action. While specialized cytoskeletal stabilization buffers (such as PHEM or CBS, which incorporate physiological Mg^2+^ and EGTA to stabilize actin networks, microtubules, and focal adhesions are often used during fluid swaps, standard DPBS was chosen here to ensure a broadly accessible, standardized workflow^32,33^. Distinguishing the relative contributions of these mechanisms, for example through time-resolved traction imaging during fixation or fixation under tonicity-matched conditions, can be possibly be investigated in future work^13^. Since PFA fixation leaves spatial displacement patterns and vector directions remain relatively intact we suggest that focal adhesion sites cross-link quickly enough to anchor the cell’s spatial footprint to the substrate, even as internal tension-generation drops^10,11,34^. Clearly, fixation artifacts may vary for different biological systems. We also mention that it is possible that quantified fixation-induced perturbations are slightly compounded by the inverse nature of traction-force reconstruction. TFM is a model-dependent inference problem in which uncertainty is affected by the inverse problem in addition to the experimental precision^35–37,5^. However, the results clearly show the importance of carefully selecting fixation protocols when quantitative force measurements are required.

Strain energy was seen to be a robust estimate of mechanical activity. Although consistently underestimated after fixation, it remains strongly correlated with live-cell values and preserves relative differences between samples. The integrated nature of strain energy, which reduces the impact of local fluctuations, directional errors, and spatial variability, renders it a more stable measure of global cellular mechanical output than traction-force magnitude.

Overall, this work demonstrates that classical TFM methods can be reliably adapted for use on fixed cells, particularly when focusing on spatial force distribution and integrated quantities such as strain energy. The developed multi-well workflow provides a straightforward method for standardized acquisition of fixed-cell TFM data.

## Materials and Methods

### Cell Culture

Human foreskin fibroblasts (ATCC; SCRC-1041) were cultured at 37 °C and in 5% CO2 in Dulbecco’s modified Eagle’s medium (DMEM; Sigma-Aldrich, D6546), supplemented with 10% fetal bovine serum (FBS, Sigma-Aldrich, F2442), 4 mM L-glutamine (Sigma-Aldrich, G7513), and 20 μg/mL gentamicin (Sigma-Aldrich, G1397). Cells were split every 2 days at a 1:6 ratio.

### PDMS Substrate Preparation for TFM

Glass coverslips (Marienfeld, 60 × 24 mm, #1.5H) were thoroughly cleaned in two steps. Initially, they were sonicated in a 5% v/v Hellmanex III solution for 10 min, followed by sonication in 100% ethanol. During each step, the coverslips were rigorously washed in double-distilled water for 5 min. Finally, the coverslips were dried under a nitrogen flow.

Silicone substrates were prepared following previous studies, using a 9:10 or 1:1 weight ratio of the two components, Cy-A and Cy-B (Dow DOWSIL CY 52-276 Kit) resulting in substrate Young’s modulus of approximately 3 or 12 kPa. The solution was mixed thoroughly, centrifuged, and degassed before 500 μL of it was pipetted onto a glass coverslip. The solution was spin-coated for 90 s using a homemade spinner, with acceleration and deceleration occurring every 30 s, to create a smooth, uniform surface. The substrates were cured at 70 °C for 30 min and were stored at room temperature (RT). For silanization of the silicone surface, the substrates were treated with a solution of 10% (v/v) 3-aminopropyl trimethoxysilane (APTES) in pure ethanol for 1.5 h. The elastomers were washed with pure ethanol, a 1:1 volume ratio of ethanol and PBS, and finally with PBS alone, each for three cycles.

As a cell culture and imaging chamber, an eight-well bottomless sticky-slide (Ibidi, 80828) was mounted on top of each PDMS substrate. After washing the chambers three times with PBS (Sigma-Aldrich, D8537), a solution of 100 nm, red-orange fluorescent (565/580) FluoSpheres carboxylate-modified microspheres (Invitrogen, F8794) at a dilution of 1:1,000 in PBS and a final concentration of 100 μg/mL 1-Ethyl-3-(3-Dimethylaminopropyl)carbodiimid, Hydrochlorid (EDC, Invitrogen, E2247) was pipetted onto the top surface of the elastomeric substrate. The solution was incubated for 20 min at RT and washed three times with PBS. Next, the substrates were coated with a solution of 10 μg/mL fibronectin (Sigma-Aldrich, F1141) with 100 μg/mL of EDC in PBS and incubated for 2 h at RT. After incubation, the chambers underwent multiple washes, first with PBS and then with cell culture medium, before being seeded with cells.

### Traction Force Measurements

To investigate differences in traction forces exerted by live and fixed fibroblasts, reference images of fluorescent beads were first acquired. Imaging was performed at 40× magnification (NA = 0.75, PH) using an inverted Olympus IX83 microscope. In each well, nine different regions were selected. Each region consisted of a 3 × 3 tiled image covering an area of approximately 650 × 650 µm². The coordinates of each region were saved using the motorized stage to enable repeated imaging of the same fields of view. Reference images were acquired prior to cell seeding to avoid the need for a cell-detachment step after fixation. After acquisition of the reference images, fibroblasts were seeded at a density of 5,000 cells/cm² in each well and incubated for 4 h under standard culture conditions. Following incubation, the DMEM medium was replaced with PBS, and the previously recorded regions were re-imaged. Images of the bead channel and corresponding bright-field images of live cells were acquired. Immediately after live-cell imaging, cells were fixed. To minimize mechanical disturbances during fixation, 200 µL of PBS was maintained in each well while 200 µL of 2× concentrated fixative solution (dissolved or diluted in PBS) was added directly to achieve final concentrations of either 4% paraformaldehyde (PFA, Sigma-Aldrich, P6148), 0.5% glutaraldehyde (GA, Sigma-Aldrich, G5882), or a mixture of 4% PFA and 0.5% GA. Cells were fixed for 20 min at room temperature (RT). Subsequently, wells were washed three times with PBS, and the same fields of view were imaged again in both the bead and bright-field channels. To prevent thermal expansion, thermal contraction, or mechanical drift artifacts associated with temperature transitions, all imaging phases, including stress-free reference substrate acquisition, live-cell imaging, and post-fixation imaging, were conducted at RT.

### Immunostaining

For immunostaining, cells were permeabilized with 0.1% Triton X-100 (Sigma-Aldrich) in PBS for 10 min at room temperature (RT), followed by three washes with PBS. Samples were then blocked with 5% bovine serum albumin (BSA, Sigma-Aldrich, A7030) in PBS for 30 min at RT. After blocking, cells were incubated with a mouse anti-paxillin primary antibody (BD Biosciences, 612405) diluted 1:500 in blocking buffer for 1 h at RT. Cells were washed three times with PBS and subsequently incubated with an Alexa Fluor 488 donkey anti-mouse IgG secondary antibody (Invitrogen, A21202) diluted 1:1000 in PBS for 1 h at RT. Following three additional PBS washes, F-actin was counterstained by incubating cells with Alexa Fluor 647–phalloidin (ThermoFisher Scientific, A22287) diluted 1:300 in blocking buffer for 30 min at RT. Finally, samples were washed three times with PBS and imaged.

### Data Analysis

Image stacks containing fluorescent bead images acquired under reference (stress-free), live-cell, and fixed-cell conditions were stacked and aligned to correct for stage drift prior to analysis. Processing and force reconstruction were performed in MATLAB (Version R2020b) using customized scripts. The entire analysis workflow includes bead detection and tracking, traction field reconstruction, maximum traction, and strain energy statistical analysis. Substrate displacement fields were computed by tracking fluorescent bead positions between live/fixed states and the reference frame. Experimental stage drift and global motion were corrected using bead positions in cell-free background regions devoid of cells. These cell-free regions also served as an internal control for intrinsic substrate stability; the absence of uncorrected background displacement confirmed that the PDMS substrate experienced no detectable viscoelastic relaxation over the 20-minute fixation period.

Prior to feature detection, all bead images were pre-processed with a Difference-of-Gaussians (DoG) bandpass filter defined by *I*_DoG_ = *G*_σ1_ ∗ *I* − *G*_σ2_ ∗ *I*, where *I* is the raw image and *G*_σ_ is a two-dimensional Gaussian kernel. Standard deviations of *σ*_1_ = 1.0 pixel (0.16 *μ*m) and *σ*_2_ = 2.0 pixels (0.33 *μ*m) were applied to all images to suppress pixel noise and slowly varying background intensity gradients while enhancing bead-sized features. Fluorescent bead feature points were detected using the MATLAB function detectMinEigenFeatures (Computer Vision Toolbox), which implements the Shi– Tomasi minimum-eigenvalue corner detector. Bead displacements were tracked using pyramidal Kanade–Lucas–Tomasi (KLT) optical flow, implemented via the MATLAB vision.PointTracker System object^38^. The tracking neighborhood window was set via the BlockSize property to [31,31] pixels, which contained approximately 13–20 tracked beads at the experimental density. This size made the gradient matrix well-conditioned and robust against image noise while defining the effective spatial resolution limit of displacement tracking. Three pyramid levels (NumPyramidLevels = 3) were used to track large displacements, and candidate points exceeding a forward–backward error threshold of 1 pixel (MaxBidirectionalError = 1) were discarded. Tracked displacements were interpolated onto a regular grid with a spacing of 7 pixels (1.14 *μ*m) using a calibrated pixel size of 0.1625 *μ*m/px.

Traction forces were reconstructed from displacement fields using Regularized Fourier Transform Traction Cytometry (FTTC) with a Poisson’s ratio of *ν* = 0.5^35^. Fixed regularization parameters were chosen for each substrate stiffness rather than using per-image automated or Bayesian selection. While Bayesian methods optimize regularization parameter based on noise characteristics, they rely on user-defined priors that introduce operator variability. Furthermore, because fixation can alter background noise levels, automated per-image optimization risks assigning different regularization parameter values to live and fixed images of the same cell, introducing mathematical artifacts into paired force comparisons. Fixing regularization parameter per stiffness ensures an identical inverse transformation across conditions, so that measured force changes reflect true physical differences in substrate displacement^35^. Regularization was scaled according to substrate Young’s modulus (*E*) and spatial resolution, yielding effective square-root Tikhonov parameters 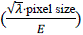 of 0.0012 *μ*m/Pa for 3 kPa substrates and 0.0003 *μ*m/Pa for 12 kPa substrates.

To quantitatively evaluate live-versus fixed-cell fields for a single cell, paired displacement vectors **u***_i_* = [*u_x_*_,*i*_, *u_y_*_,*i*_] and traction vectors **t***_i_* = [*t_x_*_,*i*_, *t_y_*_,*i*_] were evaluated at grid points *i* = 1, …, *N* within the analyzed field of view, where subscripts L and F denote live and fixed states, respectively. Local vector magnitudes were calculated as the Euclidean norm 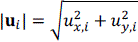, and the mean displacement magnitude (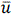) across *N* grid points was calculated as 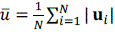, with mean traction magnitude 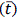 defined analogously. Metric preservation across fixation was quantified through retention metrics. Traction retention (*R*_trac_) was defined as the ratio of the mean traction magnitudes, 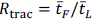, and strain energy retention (*R*_energy_) was defined as the ratio of integrated substrate strain energies (*R*_energy_ = *U*_F_/*U*_L_), where the total strain energy is given by 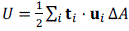 with Δ*A* implying the unit area of each grid. Local directional similarity was quantified by computing the cosine of the angle *θ_i_* between corresponding live and fixed vectors as cos *θ_i_* = S**u**_L,*i*_ ⋅ **u**_F,*i*_T/S|**u**_L,*i*_||**u**_F,*i*_|T, and averaging cos *θ_i_* over active grid points *S* to obtain the mean directional cosine 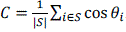. For displacements, *S* comprised points with *u*_L,*i*_ > *P*_20_, whereas for tractions, *S* included grid points where *t*_L,*i*_ exceeded 20% of the maximum live-cell traction magnitude (*t*_L,*i*_ > 0.2 ⋅ *t*_max,L_). Overall spatial pattern preservation was evaluated using Pearson’s correlation coefficient (*r*_spatial_) calculated across concatenated spatial vector components **U** = [*u_x_*_,1_, …, *u_x_*_,*N*_, *u_y_*_,1_, …, *u_y_*_,*N*_] (comprising 2*N* entries) between live and fixed states:

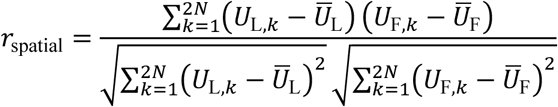

where 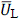 and 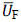 represent scalar means of the concatenated component vectors. Concatenating *x* and *y* components ensures that *r*_spatial_ captures both spatial field layout and vector directionality simultaneously, whereas for tractions, spatial correlation was evaluated across scalar traction magnitudes *t_i_*.

Independent wells/preparations were treated as the experimental units, while individual cells within each well were used to calculate well-level summary measurements and were not treated as independent replicates. When a significant main effect or interaction was detected, pairwise post-hoc comparisons were performed with correction for multiple comparisons. Because sample size was *n* = 3 wells per group, formal diagnostics for normality and homoscedasticity had limited power; conservative Tukey-Kramer corrections were applied for post-hoc comparisons. For visualization of overall cell-level relationships, scatter plots (e.g., live-vs-fixed displacement, traction force, and strain energy) depict pooled individual cells alongside Pearson correlation coefficients (*r*) and linear regression trends; however, formal hypothesis testing and group comparisons were conducted exclusively at the well level as described above. Statistical significance was defined as *P* < 0.05. Unless otherwise stated, data are presented as mean ± standard deviation (SD).

## Acknowledgments

AM and BS received funding from the European Research Council (ERC) under the European Union’s Horizon 2020 research and innovation program (BacForce, G.A.No. 852585) and from the Deutsche Forschungsgemeinschaft (DFG, German Research Foundation), Project no 492014049.

## Notes

### Competing Interest Statement

The authors have declared no competing interest.

## References

1. DuFort, C. C., Paszek, M. J. & Weaver, V. M. Balancing forces: architectural control of mechanotransduction. Nat. Rev. Mol. Cell Biol. 12, 308–319 (2011).

2. Jansen, K. A. et al. A guide to mechanobiology: Where biology and physics meet. Biochim. Biophys. Acta 1853, 3043–3052 (2015).

3. Polacheck, W. J. & Chen, C. S. Measuring cell-generated forces: a guide to the available tools. Nat. Methods 13, 415–423 (2016).

4. Roca-Cusachs, P., Conte, V. & Trepat, X. Quantifying forces in cell biology. Nat. Cell Biol. 19, 742–751 (2017).

5. Sabass, B., Gardel, M. L., Waterman, C. M. & Schwarz, U. S. High Resolution Traction Force Microscopy Based on Experimental and Computational Advances. Biophys. J. 94, 207–220 (2008).

6. Zancla, A., Mozetic, P., Orsini, M., Forte, G. & Rainer, A. A primer to traction force microscopy. J. Biol. Chem. 298, 101867 (2022).

7. Cole, R. Live-cell imaging: The cell’s perspective. Cell Adhes. Migr. 8, 452–459 (2014).

8. Sabatini, D. D., Bensch, K. & Barrnett, R. J. Cytochemistry and electron microscopy. The preservation of cellular ultrastructure and enzymatic activity by aldehyde fixation. J. Cell Biol. 17, 19– 58 (1963).

9. Karnovsky, M. L. A formaldehyde-glutaraldehyde fixative of high osmolality for use in electron-microscopy. J. Cell Biol. (1965).

10. Hopwood, D. Fixatives and fixation: a review. Histochem. J. 1, 323–360 (1969).

11. Kiernan, J. A. Formaldehyde, Formalin, Paraformaldehyde And Glutaraldehyde: What They Are And What They Do. Microsc. Today 8, 8–13 (2000).

12. Hermanson, G. T. Chapter 1 – Introduction to Bioconjugation. in Bioconjugate Techniques (Third Edition) (ed. Hermanson, G. T.) 1–125 (Academic Press, Boston, 2013). doi:10.1016/B978-0-12-382239-0.00001-7.

13. Bahr, G. F., Bloom, G. & Friberg, U. Volume changes of tissues in physiological fluids during fixation in osmium tetroxide or formaldehyde and during subsequent treatment. Exp. Cell Res. 12, 342–355 (1957).

14. Melan, M. A. & Sluder, G. Redistribution and differential extraction of soluble proteins in permeabilized cultured cells Implications for immunofluorescence microscopy. J. Cell Sci. 101, 731– 743 (1992).

15. Small, J. V., Herzog, M. & Anderson, K. Actin filament organization in the fish keratocyte lamellipodium. J. Cell Biol. 129, 1275–1286 (1995).

16. Whelan, D. R. & Bell, T. D. M. Image artifacts in Single Molecule Localization Microscopy: why optimization of sample preparation protocols matters. Sci. Rep. 5, 7924 (2015).

17. Hobro, A. J. & Smith, N. I. An evaluation of fixation methods: Spatial and compositional cellular changes observed by Raman imaging. Vib. Spectrosc. 91, 31–45 (2017).

18. Braet, F., Rotsch, C., Wisse, E. & Radmacher, M. Comparison of fixed and living liver endothelial cells by atomic force microscopy. Appl. Phys. A 66, S575–S578 (1998).

19. Yamane, Y. et al. Quantitative analyses of topography and elasticity of living and fixed astrocytes. J. Electron Microsc. (Tokyo) 49, 463–471 (2000).

20. Kim, S.-O., Kim, J., Okajima, T. & Cho, N.-J. Mechanical properties of paraformaldehyde-treated individual cells investigated by atomic force microscopy and scanning ion conductance microscopy. Nano Converg. 4, 5 (2017).

21. Zemljič Jokhadar, Š., et al. Cortical stiffness of keratinocytes measured by lateral indentation with optical tweezers. PLoS ONE 15, e0231606 (2020).

22. Li, X. et al. A high throughput microfluidic system with large ranges of applied pressures for measuring the mechanical properties of single fixed cells and differentiated cells. Biomicrofluidics 16, 034102 (2022).

23. Bergert, M. et al. Confocal reference free traction force microscopy. Nat. Commun. 7, 12814 (2016).

24. Ghagre, A. et al. Pattern-Based Contractility Screening, a Reference-Free Alternative to Traction Force Microscopy Methodology. ACS Appl. Mater. Interfaces 13, 19726–19735 (2021).

25. Beussman, K. M. et al. Black Dots: High-Yield Traction Force Microscopy Reveals Structural Factors Contributing to Platelet Forces. Acta Biomater. 163, 302–311 (2023).

26. Yang, Y. et al. Revelation of adhesive proteins affecting cellular contractility through reference-free traction force microscopy. J. Mater. Chem. B 12, 3249–3261 (2024).

27. Teo, J. L., Lim, C. T., Yap, A. S. & Saw, T. B. A Biologist’s Guide to Traction Force Microscopy Using Polydimethylsiloxane Substrate for Two-Dimensional Cell Cultures. STAR Protoc. 1, 100098 (2020).

28. Huang, B., Bates, M. & Zhuang, X. Super-Resolution Fluorescence Microscopy. Annu. Rev. Biochem. 78, 993–1016 (2009).

29. Lelek, M. et al. Single-molecule localization microscopy. Nat. Rev. Methods Primer 1, 39 (2021).

30. Ichikawa, T. et al. Chemical fixation creates nanoscale clusters on the cell surface by aggregating membrane proteins. *Commun*. Biol. 5, 487 (2022).

31. Zhao, S. et al. Fixation-induced cell blebbing on spread cells inversely correlates with phosphatidylinositol 4,5-bisphosphate level in the plasma membrane. FEBS Open Bio 4, 190–199 (2014).

32. Zhang, K. & Chen, J. The regulation of integrin function by divalent cations. Cell Adhes. Migr. 6, 20–29 (2012).

33. Leyton-Puig, D. et al. PFA fixation enables artifact-free super-resolution imaging of the actin cytoskeleton and associated proteins. Biol. Open 5, 1001–1009 (2016).

34. Stricker, J., Falzone, T. & Gardel, M. L. Mechanics of the F-actin cytoskeleton. J. Biomech. 43, 9–14 (2010).

35. Huang, Y. et al. Traction force microscopy with optimized regularization and automated Bayesian parameter selection for comparing cells. Sci. Rep. 9, 539 (2019).

36. Blumberg, J. W. & Schwarz, U. S. Comparison of direct and inverse methods for 2.5D traction force microscopy. PLOS ONE 17, e0262773 (2022).

37. Kandasamy, A., Yeh, Y.-T., Serrano, R., Mercola, M. & del Alamo, J. C. Uncertainty-aware traction force microscopy. PLOS Comput. Biol. 21, e1013079 (2025).

39. Holenstein, C. N., Silvan, U. & Snedeker, J. G. High-resolution traction force microscopy on small focal adhesions – improved accuracy through optimal marker distribution and optical flow tracking. Sci. Rep. 7, 41633 (2017).

