## Supplementary material for "Mechanical Fidelity of Fixed-Cell Traction Force Microscopy": Supp. Fig. 1 and Supp. Tab. 1

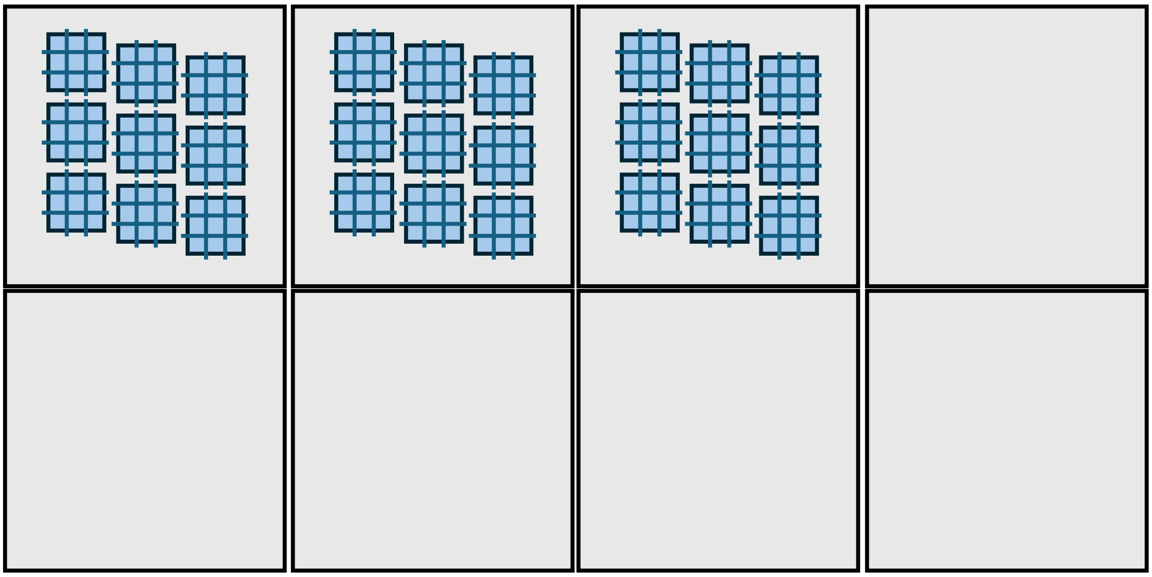

**Supplementary Fig. 1**. Schematic representation of imaging field of views.

**Supplementary Table 1**. Two-Way ANOVA and Post-Hoc Tukey-Kramer Comparisons

| **Analysis** | **Metric** | **Factor / Comparison** | **DF Effect, Error** | **F-value** | **Mean Difference (%)** | **p-value** | **Significance** |
|---|---|---|---|---|---|---|---|
| **Two-Way ANOVA** | **Displacement Retention** | Fixation Method | 2, 12 | 5.78 | — | 0.0185 | **Significant (*)** |
| | | Substrate Stiffness | 1, 12 | 0.04 | — | 0.8541 | Not Significant |
| | | Method × Stiffness | 2, 12 | 0.08 | — | 0.9210 | Not Significant |
| | **Force Retention** | Fixation Method | 2, 12 | 6.02 | — | 0.0169 | **Significant (*)** |
| | | Substrate Stiffness | 1, 12 | 0.01 | — | 0.9231 | Not Significant |
| | | Method × Stiffness | 2, 12 | 0.06 | — | 0.9412 | Not Significant |
| | **Strain Energy Retention** | Fixation Method | 2, 12 | 5.41 | — | 0.0221 | **Significant (*)** |
| | | Substrate Stiffness | 1, 12 | 0.02 | — | 0.8890 | Not Significant |
| | | Method × Stiffness | 2, 12 | 0.07 | — | 0.9324 | Not Significant |
| | **Directional Cosine** | Fixation Method | 2, 12 | 0.89 | — | 0.4352 | Not Significant |
| | | Substrate Stiffness | 1, 12 | 1.15 | — | 0.3045 | Not Significant |

| Analysis | Metric | Factor / Comparison | DF Effect, Error | F-value | Mean Difference (%) | p-value | Significance |
|---|---|---|---|---|---|---|---|
| | | Method × Stiffness | 2, 12 | 0.64 | — | 0.5401 | Not Significant |
| | **Spatial Correlation (r)** | Fixation Method | 2, 12 | 0.18 | — | 0.8389 | Not Significant |
| | | Substrate Stiffness | 1, 12 | 0.45 | — | 0.5152 | Not Significant |
| | | Method × Stiffness | 2, 12 | 0.21 | — | 0.8134 | Not Significant |
| **Post-Hoc Tukey-Kramer** | **Displacement Retention** | GA vs. PFA | — | — | **+24.8** | 0.0152 | **Significant** |
| | | GA vs. Mix | — | — | +14.0 | 0.1842 | Not Significant |
| | | Mix vs. PFA | — | — | +10.8 | 0.3210 | Not Significant |
| | **Force Retention** | GA vs. PFA | — | — | **+14.5** | 0.0134 | **Significant** |
| | | GA vs. Mix | — | — | +8.3 | 0.2140 | Not Significant |
| | | Mix vs. PFA | — | — | +6.2 | 0.3812 | Not Significant |
| | **Strain Energy Retention** | GA vs. PFA | — | — | **+24.2** | 0.0224 | **Significant** |

| Analysis | Metric | Factor Comparison | DF Effect, / Error | F-value | Mean Difference (%) | p-value | Significance |
|---|---|---|---|---|---|---|---|
| | | GA vs. Mix | — | — | +13.1 | 0.2205 | Not Significant |
| | | Mix vs. PFA | — | — | +11.1 | 0.3150 | Not Significant |

*Statistical summary of two-way ANOVA models evaluated at the well level (n = 3 independent wells per condition; total N = 18 wells, summarizing N = 267 total single cells). Significant main effects of Fixation Method were followed by Tukey-Kramer pairwise comparisons. α = 0.05. GA, glutaraldehyde; PFA, paraformaldehyde; Mix, mixed fixation method. *p < 0.05.*
